# Swimming performance traits of twenty-one Australian fish species: a fish passage management tool for use in modified freshwater systems

**DOI:** 10.1101/861898

**Authors:** Jabin R. Watson, Harriet R. Goodrich, Rebecca L. Cramp, Matthew A. Gordos, Yulian Yan, Patrick J. Ward, Craig E. Franklin

## Abstract

Freshwater ecosystems have been severely fragmented by artificial in-stream structures designed to manage water for human use. Significant efforts have been made to reconnect freshwater systems for fish movement, through the design and installation of dedicated fish passage structures (fishways) and by incorporating fish-sensitive design features into conventional infrastructure (e.g. culverts). Key to the success of these structures is making sure that the water velocities within them do not exceed the swimming capacities of the local fish species. Swimming performance data is scarce for Australian fish, which have a reduced swimming capacity when compared to many North American and European species. To help close this knowledge gap and assist fisheries management and civil engineering, we report the swimming performance capacities of twenty-one small-bodied fish and juveniles (< 10 cm) of large bodied species native to Australia as measured by critical swimming speed (*U*crit) and burst swimming speed (*U*sprint) in a recirculating flume. This data is complemented by endurance swim trials in a 12-meter hydraulic flume channel, and by measures of flume traverse success. Building on the utility of this dataset, we used a panel of morphological, behavioural and ecological traits to first assess their relative contributions to the observed swimming performance data, and second, to determine if they could be used to predict swimming performance capacity – a useful tool to assist in the management of species of conservation concern where access to swimming performance data may be limited. We found that body length combined with depth station (benthic, pelagic or surface) explained most of the interspecific variation in observed swimming performance data, followed by body shape and tail shape. These three traits were the most effective at predicting swimming performance in a model/unknown fish. This data will assist civil engineers and fisheries managers in Australia to mitigate the impact of in-stream structures on local fish populations.

## Introduction

Fish are a major component of freshwater ecosystems, with healthy fish populations reflective of a healthy system (Durance et al., 2016; Lynch et al., 2016; O’Brien et al., 2016). Globally, freshwater fish populations have been severely impacted by human activities including direct over-exploitation, pollution, habitat degradation, and habitat fragmentation which has resulted in severe reductions and, in certain cases, extirpation of fish populations and species (Arthington et al., 2016; Grill et al., 2019). A significant cause of freshwater habitat fragmentation is the construction of civil instream structures like dams, weirs and culverts which block native fish migration. Historically these structures were designed and built with little to no consideration of their impact on the surrounding aquatic communities. However, their deleterious impact on fish populations is now well acknowledged, with extensive remediation efforts underway globally to reconnect fragmented freshwater systems through the incorporation of dedicated design elements that facilitate fish passage (Gordon et al., 2018; Harris et al., 2016; Mao, 2018; Noonan et al., 2012; Katopodis and Williams, 2012; Roscoe and Hinch, 2010).

A multitude of fish-passage designs exist, reflective of the varied biological requirements found across the diversity of freshwater fish fauna. Unsurprisingly, fish passage structures have evolved in an iterative fashion over the last century, building upon both successes and failures (Katopodis and Williams, 2012). Selecting a specific fish passage design for a particular barrier is often complicated by the size of the barrier, site-specific constraints such as hydrology and gradient, and the ecology, behaviour and physiology of the fish that need to be passed (Lapointe et al., 2013; Palmer et al., 2007). The implementation of a particular design at a fish passage barrier is a complex task, requiring the balancing of civil requirements with that of the local fish species while operating within budgetary constraints (Hyde, 2015). Ultimately the biology needs to drive the design of ‘fish-friendly’ structures. The greater our understanding of the fish species requiring passage, the higher the probability that the fish passage structure will function successfully.

Fish swimming performance is a key biological aspect of fish passage structure design (Botha et al., 2018; Cai et al., 2018; Scruton et al., 1998; Bice and Zampatti, 2005; Katopodis et al., 2019; Link et al., 2017; Peake, 2014; Starrs et al., 2011). Excessive water velocities and turbulence are major physical barriers to fish movement that are often generated by concentrated flows associated with instream structures. Empirical experimental fish swimming performance studies can provide important baseline data that can be used by managers to guide the design of fish passage structures, or to provide evidence-backed advice on appropriate water use practices to limit impacts on fish populations (Cooke et al., 2017). Additionally, carefully designed experiments conducted in controlled conditions can test cause-and-effect relationships, which can be difficult to elucidate in the field due to inherent complexities (Cooke et al., 2017).

Fish swim using different gaits that can be classified into three categories largely dependent on the duration of time that the swimming speed is maintained, and subsequently, the type of muscles engaged (Beamish, 1978). Sustained swimming speeds generally involve durations longer than 200 minutes where red muscle, powered by aerobic metabolism, can work with limited accumulation of metabolic waste products that induce muscle fatigue. Burst swimming speeds can be maintained for less than 20 seconds due to its reliance on white muscle. White muscle has low aerobic capacity resulting in the rapid depletion of energy stores and the accumulation of metabolic waste products within muscle cells causing fatigue. Prolonged swimming speeds (*U*crit) span the gap between burst and sustained, utilising both types of muscle and energy production pathways (Beamish, 1978). Typically, prolonged and burst swimming gaits are required for fish to successfully traverse upstream of man-made barriers. In addition, structure traversability estimates provide useful information on the ability of a fish to cover ground against the direction of water flow, using whichever mode of swimming they choose. Traversability success indices record the capacity of fish to move a predefined distance in a swimming environment, usually against the flow of water. Laboratory based performance tests enable the hydraulic environment to be simplified and adjusted in a consistent and controllable manner in order to assess the impacts of a particular feature on the capacity of fish to move through the environment. Absolute performance in a fish passage structure is likely to be highly context-specific, since animals may choose to use a variety of gaits and behaviours to traverse a structure dependent on the specific range of biotic and abiotic conditions in and around the structure. The use of multiple performance tests that incorporate both volitional and non-volitional measures provides a more holistic assessment of the performance range of a species. Ultimately, for swimming performance measures to be a useful management tool in mixed-species communities, they need to be quantified using the same protocol and equipment since fish swimming performance is known to be highly sensitive to variations in protocol (i.e. relative velocity increases, pre-test training/acclimation, velocity increment length; Brett, 1967) and test equipment (Kern et al., 2018), as well as other performance variables (body mass, temperature, developmental stage, wild-caught vs hatchery stock, etc.) (Kern et al., 2018). Until recently, a lack of consistency across existing fish swimming performance testing and handling protocols has meant that we have had limited capacity for interspecific comparisons of fish swimming performance (Kern et al., 2018).

Swimming performance data for native Australian fish is sparse (see (Kern et al., 2017; Kopf et al., 2014; Mallen-Cooper, 1992; Rodgers et al., 2017, 2014; Starrs et al., 2011; Bice and Zampatti, 2005; Whiterod, 2012)), and what does exist has been collected using different protocols and equipment. As an aid for fisheries managers in Australia, we aimed to quantify the swimming performance of twenty-one native fish species. Small-bodied species, and juveniles of larger growing fish species are the most vulnerable to the velocity challenges caused by man-made in-stream structures due to body size-dependent swimming performance capacity. Consequently, we included fifteen small-bodied species and juveniles of six species that are large bodied as adults (see common adult sizes in Table 1.).

**Table 1.** Physical and behavioural characteristics of study species.

| Scientific name | Common name | Common adult size (mm) | Location | Preferred habitat | Migration classification | Movement distance * | Body shape | Tail shape | Depth station | Swimming mode |
| --- | --- | --- | --- | --- | --- | --- | --- | --- | --- | --- |
| *Trachystoma petardi* | Freshwater mullet | 400 | Coastal | Channel specialist | Catadromous | Macro | Fusiform | Indented | Surface | Sustained |
| *Maccullochella peelii* | Murray cod | 700 | Inland | Channel specialist | Potamodromous | Meso | Compressiform | Rounded | Pelagic | Slow Sustained / Burst |
| *Macquaria ambigua* | Golden perch | 500 | Inland | Channel specialist | Potamodromous | Macro | Compressiform | Truncate | Pelagic | Slow Sustained / Burst |
| *Macquaria novemaculeata* | Australian bass | 400 | Coastal | Channel specialist | Catadromous | Macro | Compressiform | Indented | Pelagic | Slow Sustained / Burst |
| *Bidyanus bidyanus* | Silver perch | 300 | Inland | Channel specialist | Potamodromous | Macro | Compressiform | Indented | Pelagic | Sustained / Burst |
| *Leiopotherapon unicolor* | Spangled perch | 300 | Inland | Generalist | Potamodromous | Meso | Compressiform | Truncate | Pelagic | Slow Sustained / Burst |
| *Galaxias brevipinnis* | Climbing galaxiid | 100 | Coastal | Channel specialist | Potamodromous | Meso | Sagitiform | Truncate | Pelagic | Slow Sustained / Burst |
| *Retropinna semoni* | Australian smelt | 60 | Coastal / Inland | Generalist | Potamodromous | Micro | Fusiform | Forked | Pelagic | Sustained |
| *Craterocephalus stercusmuscarum* | Fly-specked hardyhead | 55 | Inland | Generalist | Potamodromous | Meso | Fusiform | Indented | Pelagic | Sustained |
| *Pseudomugil signifier* | Pacific blue-eye | 40 | Coastal | Generalist | Amphidromous | Micro | Fusiform | Indented | Pelagic | Sustained |
| *Melanotaenia fluviatilis* | Murray river rainbowfish | 85 | Inland | Generalist | Potamodromous | Micro | Compressiform | Indented | Pelagic | Slow Sustained / Burst |
| *Melanotaenia duboulayi* | Crimson-spotted rainbowfish | 80 | Coastal | Generalist | Potamodromous | Micro | Compressiform | Indented | Pelagic | Slow Sustained / Burst |
| *Rhadinocentrus ornatus* | Ornate rainbowfish | 40 | Coastal | Wetland specialist | Potamodromous | Micro | Fusiform | Indented | Pelagic | Slow Sustained / Burst |
| *Ambassis ambassizii* | Olive perchlet | 60 | Coastal / Inland | Wetland specialist | Potamodromous | Micro | Compressiform | Forked | Pelagic | Slow Sustained / Burst |
| *Nannoperca australis* | Southern pygmy perch | 60 | Inland | Wetland specialist | Potamodromous | Micro | Compressiform | Rounded | Pelagic | Burst |
| *Tandanus tandanus* | Eel-tailed catfish | 450 | Coastal / Inland | Generalist | Potamodromous | Meso | Compressiform | Pointed | Benthic | Slow Sustained / Burst |
| *Hypseleotris compressa* | Empire gudgeon | 80 | Coastal | Generalist | Amphidromous | Meso | Compressiform | Rounded | Benthic | Burst |
| *Hypseleotris galii* | Firetail gudgeon | 35 | Coastal | Generalist | Potamodromous | Micro | Compressiform | Rounded | Benthic | Burst |
| *Philypnodon grandiceps* | Flathead gudgeon | 80 | Coastal / Inland | Generalist | Potamodromous | Micro | Depressiform | Rounded | Benthic | Burst |
| *Mogurnda adspersa* | Purple spotted gudgeon | 80 | Coastal / Inland | Generalist | Potamodromous | Meso | Compressiform | Rounded | Benthic | Burst |
| *Redigobius bikolanus* | Speckled goby | 40 | Coastal | Generalist | Amphidromous | Meso | Compressiform | Rounded | Benthic | Burst |

\*Adapted from Mallen-Cooper and Zampatti (2012) where micro < 100 m, meso is 100s m to 10s km, and macro is 100s km.

To enable a valid inter-specific comparison, we collected *U*crit and *U*sprint data with a consistent protocol in the same flume. This was complemented by collecting traversability and endurance swimming times for fifteen of those species, with each species swum at three different fixed water velocities in a 12-meter experimental channel. By sampling this range of swimming performance metrics, we aimed to increase the applicability of the data to different situations. Finally, we combined the swimming performance data with the physical and behavioural characteristics of each study species to determine whether this data could be used collectively to improve our ability to predict fish swimming ability. Generally, when estimating the swimming capacity of an unknown species, the prediction is done using just fish length. This dataset presented an opportunity to assess the applicability of using body length alone when predicting the swimming capacity of Australian freshwater fish, as well as if any other morphological, behavioural or ecological characteristics were able to improve the predictions. This allows fisheries managers to use species included in this study as surrogates for less common or endangered species for which no swimming performance data exists. What we present here is critical baseline data that is essential for Australian fisheries managers and engineers to design effective fish passage structures.

## Methods

### Species choices and characterisations

We chose twenty-one native Australian fish species that span a range of body sizes, age classes, body shapes, behavioural characteristics and ecological traits. The physical characteristics that species were classed into were body shape and tail shape. The behavioural characteristics included preferred habitat, migration classification, movement distance (based on Mallen-Cooper and Zampatti, 2015), depth station, and swimming mode. We also included each species’ geographical location as either coastal, inland or both. Coastal indicates river systems that drain into the Pacific Ocean, and inland refers to the Murray-Darling Basin. For details see Table 1.

### Fish sources and husbandry

*M. novemaculeata, T. tandanus, H. compressa, H. galii, C. stercusmuscarum, M. petardii, M. ambigua, M. peelii, M. fluviatilis, A. ambassizii, R. ornatus, M. adspersa, B. bidyanus, N. australis* and *L. unicolor* were sourced from commercial hatcheries. *R. semoni, M. duboulayi* and *P. signifier* were collected at Moggil Creek and Cedar Creek, Brisbane (21°30’14”S 152°55’49”E and 27°19’28”S 152°47’40”E). *P. grandiceps* and *R. bikolanus* were collected at North Pine Dam, Brisbane (27°16’57”S 152°55’39”E). All collections were done using box traps under Queensland Fisheries General Fisheries Permit number 186295.

Fish were housed only with conspecifics in 40 L glass aquaria that were part of 1000 L recirculating systems. Each system had mechanical and biological filtration, and UV sterilisation. Approximately 25% of the systems water volume was exchanged per day using a constant flow of carbon filtered tap water. Water temperature was controlled at 25 ± 1 °C, with a 12:12h light-dark lighting regime being provided. The fish were fed daily to satiation using an aquaculture crumble diet (Ridley, Brisbane, Australia) and frozen bloodworms (Chironomidae). All fish were fasted for 24 hours before each performance trial to ensure a post-absorptive state (Norin et al., 2014), with individuals that had been swum being housed separately from fish that had not been tested.

### Swimming performance - Ucrit and Usprint

We quantified *U*crit values (Brett, 1964; Kern et al., 2017; Rodgers et al., 2014) for twenty native Australian fish species listed in Table 2. Likewise, we collected *U*sprint (Starrs et al., 2011) data for all species, with the exception being *G. brevipinnis*. Both *U*crit and *U*sprint trials were performed in a 185 L flow controlled recirculating flume (Loligo, Tjele, Denmark) that was calibrated using a Prandtl-pitot tube (Kern et al., 2017) and maintained at 25 ± 1 °C. For each *U*crit test, individual fish were introduced to the flume and left for five min to recuperate from handling stress. To start the trial, water velocity was set to 0.1 m s^−1^. Every 5 min, the velocity was increased by 0.05 m s^−1^ until the fish fatigued. Fatigue was defined as the fish resting on the back screen of the swimming chamber for 3 s.

**Table 2.** Summary of *U*crit and *U*sprint swimming performance data.

| Scientific name | Common name | *Ucrit* size (cm) (Mean ± SD [range]) | *Ucrit* 25<sup>th</sup> percentile (m s<sup>-1</sup>) | *Ucrit* mean (m s<sup>-1</sup>) | *Ucrit* (n) | *Usprint* size (cm) (Mean ± SD [range]) | *Usprint* 25<sup>th</sup> percentile (m s<sup>-1</sup>) | *Usprint* mean (m s<sup>-1</sup>) | *Usprint* (n) |
| --- | --- | --- | --- | --- | --- | --- | --- | --- | --- |
| *Macquaria novemaculeata* | Australian bass | 6.0 ± 0.7<br>[4.3 - 7.3] | 0.53 | 0.60 | 31 | 3.9 ± 0.5<br>[3.4 - 5.0] | 0.52 | 0.58 | 25 |
| *Retropinna semoni* | Australian smelt | 4.5 ± 0.7<br>[2.5 - 6.0] | 0.62 | 0.66 | 25 | 4.8 ± 0.7<br>[3.5 - 6.1] | 0.67 | 0.70 | 25 |
| *Melanotaenia duboulayi* | Crimson-spotted rainbowfish | 6.8 ± 1.2<br>[5.3 - 10.3] | 0.63 | 0.63 | 26 | 6.8 ± 1.2<br>[4.6 - 8.5] | 0.67 | 0.70 | 25 |
| *Tandanus tandanus* | Eel-tailed catfish | 10.0 ± 1.5<br>[6.1 - 10.7] | 0.36 | 0.41 | 42 | 8.3 ± 0.8<br>[6.9 - 9.4] | 0.46 | 0.50 | 25 |
| *Hypseleotris compressa* | Empire gudgeon | 5.3 ± 0.7<br>[4.4 - 7.8] | 0.34 | 0.44 | 23 | 5.1 ± 0.8<br>[4.1 - 6.6] | 0.49 | 0.51 | 25 |
| *Hypseleotris galii* | Firetail gudgeon | 3.8 ± 0.8<br>[2.4 - 5.4] | 0.34 | 0.38 | 25 | 3.9 ± 0.7<br>[2.5 - 5.6] | 0.48 | 0.50 | 25 |
| *Philypnodon grandiceps* | Flathead gudgeon | 4.8 ± 1.0<br>[3.2 - 6.3] | 0.18 | 0.33 | 15 | 5.3 ± 0.6<br>[3.9 - 5.8] | 0.40 | 0.47 | 13 |
| *Craterocephalus stercusmuscarum* | Fly-specked hardyhead | 4.6 ± 0.5<br>[3.7 - 5.8] | 0.46 | 0.49 | 25 | 4.8 ± 0.6<br>[3.8 - 5.9] | 0.50 | 0.55 | 25 |
| *Trachystoma petardi* | Freshwater mullet | 7.3 ± 1.0<br>[5.9 - 8.8] | 0.80 | 0.85 | 14 | 6.5 ± 1.3<br>[4.7 - 8.7] | 0.82 | 0.83 | 17 |
| *Galaxias brevipinnis* | Climbing galaxid | 9.8 ± 2.4<br>[8.1 - 11.5] | 0.40 | 0.49 | 2 | NA | NA | NA | NA |
| *Macquaria ambigua* | Golden perch | 4.7 ± 0.7<br>[3.5 - 6.4] | 0.28 | 0.32 | 25 | 5.5 ± 0.7<br>[4.2 - 7.5] | 0.43 | 0.47 | 25 |
| *Maccullochella peelii* | Murray cod | 6.7 ± 1.8<br>[4.0 - 10.0] | 0.40 | 0.46 | 43 | 6.3 ± 1.2<br>[4.3 - 8.1] | 0.57 | 0.61 | 40 |
| *Melanotaenia fluviatilis* | Murray river rainbowfish | 7.0 ± 0.6<br>[6.1 - 8.2] | 0.42 | 0.45 | 25 | 6.9 ± 0.5<br>[6.0 - 7.8] | 0.66 | 0.68 | 25 |
| *Ambassis ambassizii* | Olive perchlet | 5.8 ± 0.6<br>[4.5 - 7.0] | 0.50 | 0.53 | 24 | 5.6 ± 0.7<br>[3.5 - 6.5] | 0.49 | 0.57 | 25 |
| *Rhadinocentrus ornatus* | Ornate rainbowfish | 4.0 ± 0.9<br>[2.5 - 5.8] | 0.46 | 0.52 | 25 | 4.1 ± 0.9<br>[3.0 - 6.3] | 0.45 | 0.51 | 25 |
| *Pseudomugil signifier* | Pacific blue-eye | 4.1 ± 0.3<br>[3.5 - 4.7] | 0.42 | 0.46 | 25 | 3.5 ± 0.4<br>[2.5 - 4.0] | 0.48 | 0.52 | 25 |
| *Mogurnda adspersa* | Purple spotted gudgeon | 6.1 ± 1.8<br>[4.0 - 10.0] | 0.18 | 0.21 | 25 | 7.2 ± 2.2<br>[4.3 - 10.3] | 0.53 | 0.57 | 25 |
| *Bidyanus bidyanus* | Silver perch | 6.6 ± 0.8<br>[4.8 - 8.2] | 0.66 | 0.68 | 25 | 8. ± 0.7<br>[7.4 - 10.1] | 0.68 | 0.74 | 25 |
| *Nannoperca australis* | Southern pygmy perch | 3.9 ± 0.9<br>[2.3 - 5.6] | 0.34 | 0.35 | 25 | 4.8 ± 0.4<br>[3.8 - 5.9] | 0.44 | 0.49 | 25 |
| *Leiopotherapon unicolor* | Spangled perch | 3.6 ± 1.2<br>[2.5 - 6.7] | 0.35 | 0.41 | 13 | 6.3 ± 1.8<br>[3.0 - 9.5] | 0.61 | 0.66 | 40 |
| *Redigobius bikolanus* | Speckled goby | 3.2 ± 0.3<br>[2.6 - 3.8] | 0.32 | 0.34 | 10 | 3.0 ± 0.5<br>[2.3 - 3.8] | 0.44 | 0.43 | 11 |

For the *U*sprint tests, individual fish were transferred to the swimming chamber within the recirculating flume and left for 5 min at 0.1 m s^−1^ to orientate with the flow. After 5 min, the velocity was increased by 0.05 m s^−1^ every 10 s until the fish fatigued. After a *U*crit or *U*sprint swimming trial, fish were measured (total length) and weighed. Both *U*crit and *U*sprint were calculated as in Brett (1964):

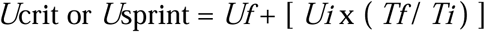

where *Uf* is the highest sustained water velocity (m s^−1^), *Ui* is the velocity increase increments (0.05 m s^−1^ for both *U*crit and *U*sprint), *Tf* is time swum during the final velocity increment, and *Ti* is the time increment (300 and 10 s for *U*crit and *U*sprint respectively).

### Swimming performance - endurance

The endurance trials were conducted in a hydraulic channel (12 × 0.5 × 0.25 m) with a constant supply header tank system (Kern et al., 2017). The channel was part of a 40,000 L recirculating fish swimming facility at The University of Queensland’s Biohydrodynamics Laboratory. The water inlet of the channel was fitted with flow straighteners, with stainless-steel screens fitted to both ends of the channel to prevent fish either entering the flow straighteners, or to catch the fish upon fatigue before the flume outflow. The system was maintained at 25 ± 1 °C with a heat pump (Oasis C58T-Vb, New Zealand) and was equipped with mechanical and biological filtration.

We quantified the swimming endurance times for fifteen species of native Australian fish, with each species being tested at three different bulk channel velocities (Table 3). The three test velocities were species-specific and determined based on the measured *U*crit swimming capacity of that species. Fish were swum individually at the pre-determined bulk channel velocity for a maximum of 60 min. If a fish did not fatigue within this 60 min time frame, that data was treated as censored or unobservable. Fatigue was defined as the fish resting on the screen at the downstream end of the channel for 3 s. We also quantified the rate of traverse success. Fish were released 1 m from the downstream end of the channel, with traverse success defined as the fish swimming up 8 m of channel to a marked point without encouragement. The 8 m distance is reflective of a dual carriageway culvert, the most common culvert size in New South Wales, Australia.

**Table 3.**
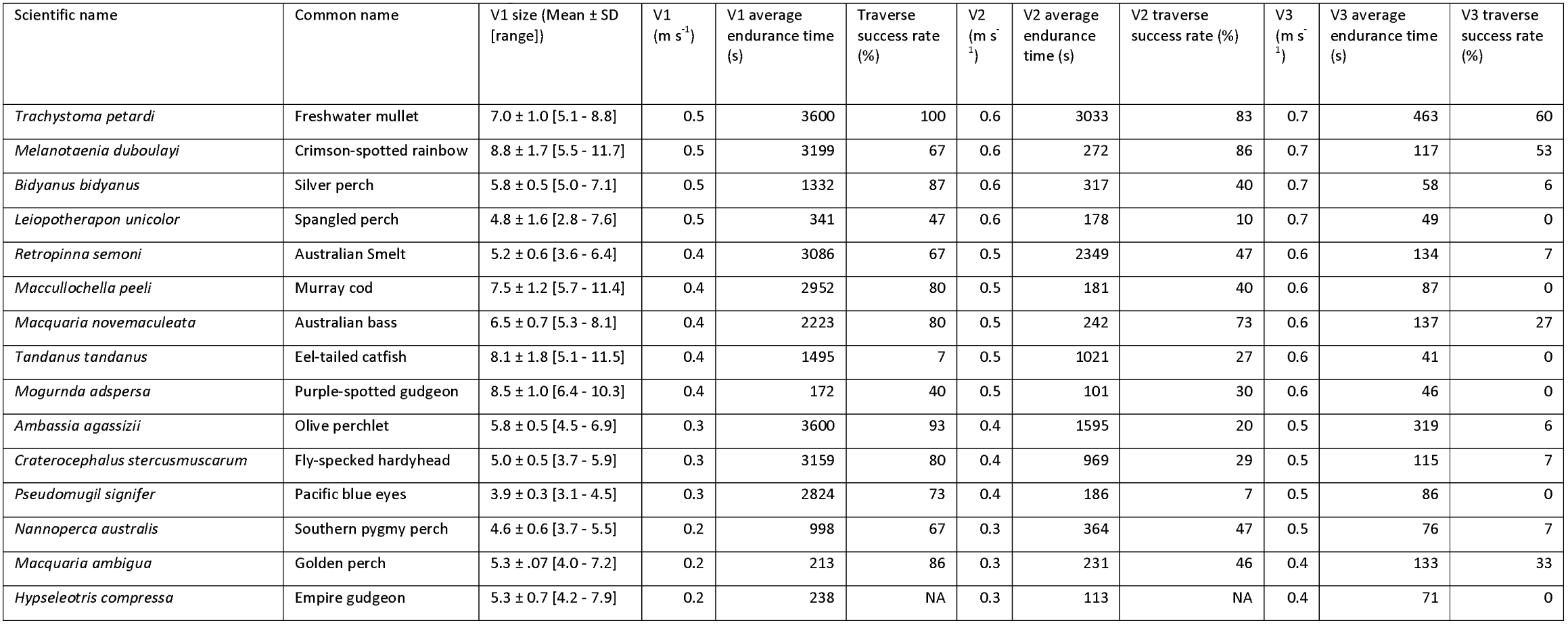
Endurance times, velocities and size ranges for fish swum in the 12 m flume channel.

### Modelled traversable distances

The 25^th^ percentile *U*crit (m s^−1^) values were calculated for each species and used to model the maximum distances a fish within the size ranges sampled may swim against a set water velocity. This information may be used to estimate the likelihood of at least 75% of fish within the size range sampled successfully traversing a structure of known length and known water velocity. This was done using the equation:

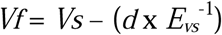

where *Vf* is the velocity in the culvert (m s^−1^) to be traversed, Vs is the 25^th^ percentile *U*crit value (m s^−1^), *d* is the length of the culvert and E_vs_^−1^ is the time increment used during the *U*crit or *U*sprint trials (s) (Peake et al., 1997). This equation takes into account that the fish must swim faster than the water velocity to achieve forward progress.

### Statistical analysis

All statistical analysis and figure generation was performed using R (R Core Team, 2018), within the *RStudio* environment (2018). Data was analysed using the following packages: lme4 (Bates et al., 2015), lsmeans (Lenth, 2016), survival (Therneau, 2015), Matrix (Bates and Maechler, 2018), Rmisc (Hope, 2013), MuMin (Bartoń, 2019), lmerTest (Kuznetsova et al., 2017) and multcomp (Hothorn et al., 2008). The following packages were used to visualise the data and generate the figures: ggplot2 (Wickham, 2016), ggfortify (Tang et al., 2016), ggpubr (Kassambara, 2018), survminer (Kassambara and Kosinski, 2018), dplyr (Wickam et al., 2018) and RColorBrewer (Neuwirth, 2014).

Mixed effects models were used to quantify how each physical and behavioural characteristic individually contributed to both *U*crit and *U*sprint. Both swimming performance metrics were analysed as m s^−1^ and BL s^−1^. The respective swimming performance metric (*U*crit or *U*sprint) was the response variable, with fish length and a physical or behavioural characteristic being treated as a fixed effect (i.e., as predictor) and species being treated as a random effect. BL s^−1^ results are reported in the supplementary information. The potential effect of each characteristic on *U*crit and *U*sprint, represented as both m s^−1^ and BL s^−1^(see supplementary information). Each physical or behavioural characteristic was modelled separately. Statistical significance was set at *p* < 0.05. First, the validity of predicting the *U*crit from body length and other morphological and ecological data was assessed by applying a linear model to the *U*crit database in an iterative fashion on one species at a time. A reference *U*crit database was then made that included all the *U*crit, length and categorical data for all species, except the one for which we were making a prediction. Next a series of linear models were generated from that reference database with *U*crit as the response variable, and each including a selection of predictors to be tested with all possible interactions. The first model included length plus all predictors (migration classification, movement distance, body shape, tail shape, depth station, swimming mode, preferred habitat and location). Mass was not included in the model as it reduced the accuracy when length was also a factor. The second model had only length as the predictor. The third, fourth and fifth models had either depth station, body shape or tail shape as a co-predictor with length. We used R’s “*predict()”* function to enter the length of the individual fish of the species for which we were predicting the data, and the respective categorisation of their species.

A mixed effect model was used to assess how the different predicted *U*crit values compared to the observed data. *U*crit was the response variable, with the predictor(s) defined by the type of model, and with species treated as a random effect. For this analysis, *P. grandiceps, T. tandanus, G. brevipinnis* and *T. petardi* were not included as they had unique characteristics (e.g. *P. grandiceps* was the only representative with a depressiform body shape), which did not allow predictive data for those characteristics.

All data is publicly available and can be found at (to be added for publication).

## Results

### Ucrit and Usprint

We analysed the data standardised to both meters and body lengths per second, with the latter reported in the supplementary information. For *U*crit or *U*sprint results for each species refer to Table 2, which also details the size ranges of fish swum. Fig. 1 and Fig. 2 compare *U*crit and *U*sprint values for all species tested, including individual data points for each fish. Overall, we found that *T. petardi* to be the best performing species with the highest average *U*crit and *U*sprint values of 0.85 m s^−1^ and 0.83 m s^−1^. Conversely the worst average *U*crit was observed by *M. adspersa* at 0.21 m s-1, and the slowest average *U*sprint was recorded by *R. bikolanus* at 0.43 m s^−1^.

**Figure 1.**
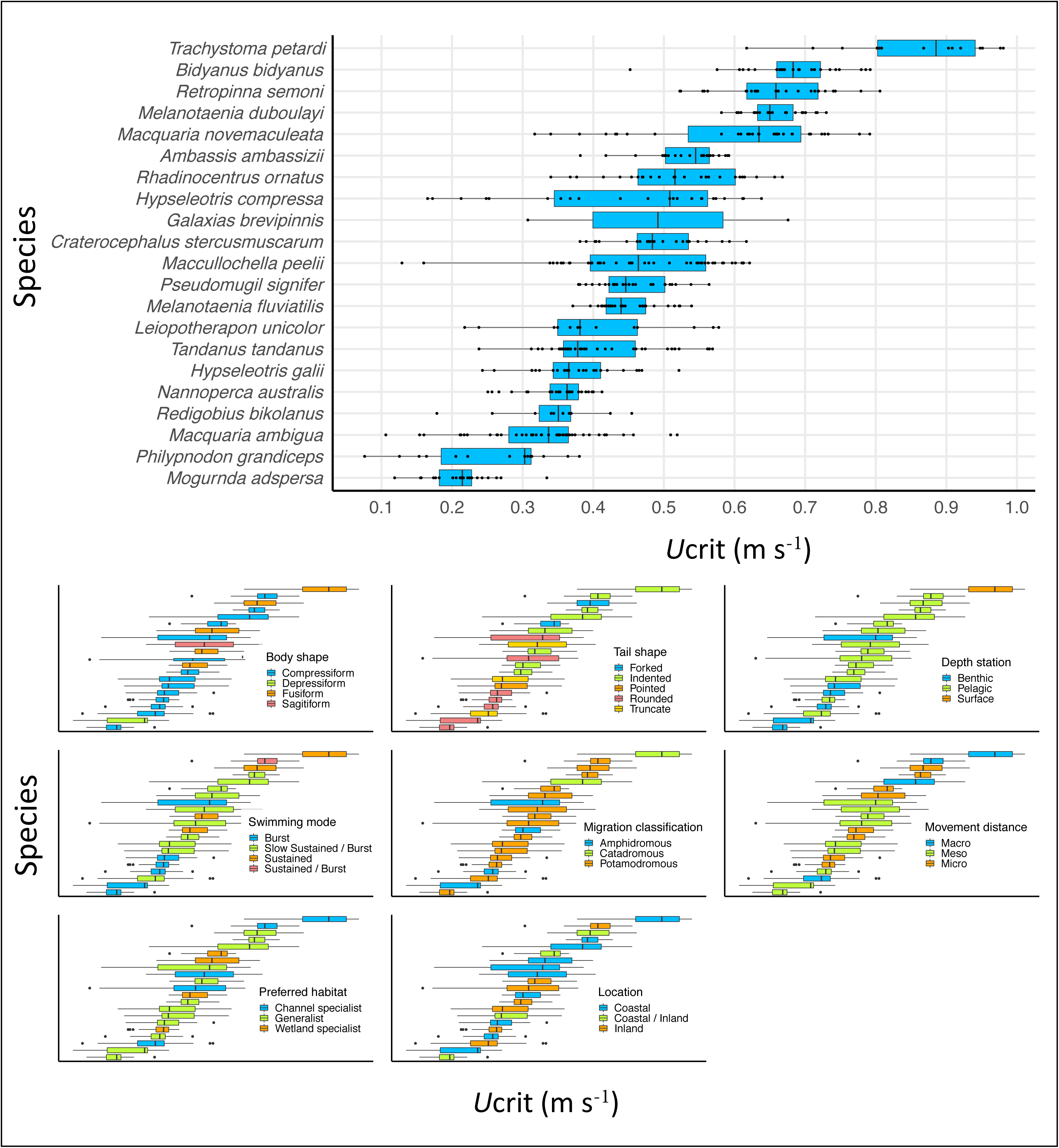
Visual comparison of the *U*crit values with individual fish shown as dots. Each of the nine physical or behavioural characteristics is plotted to show the trends across the species tested.

**Figure 2.**
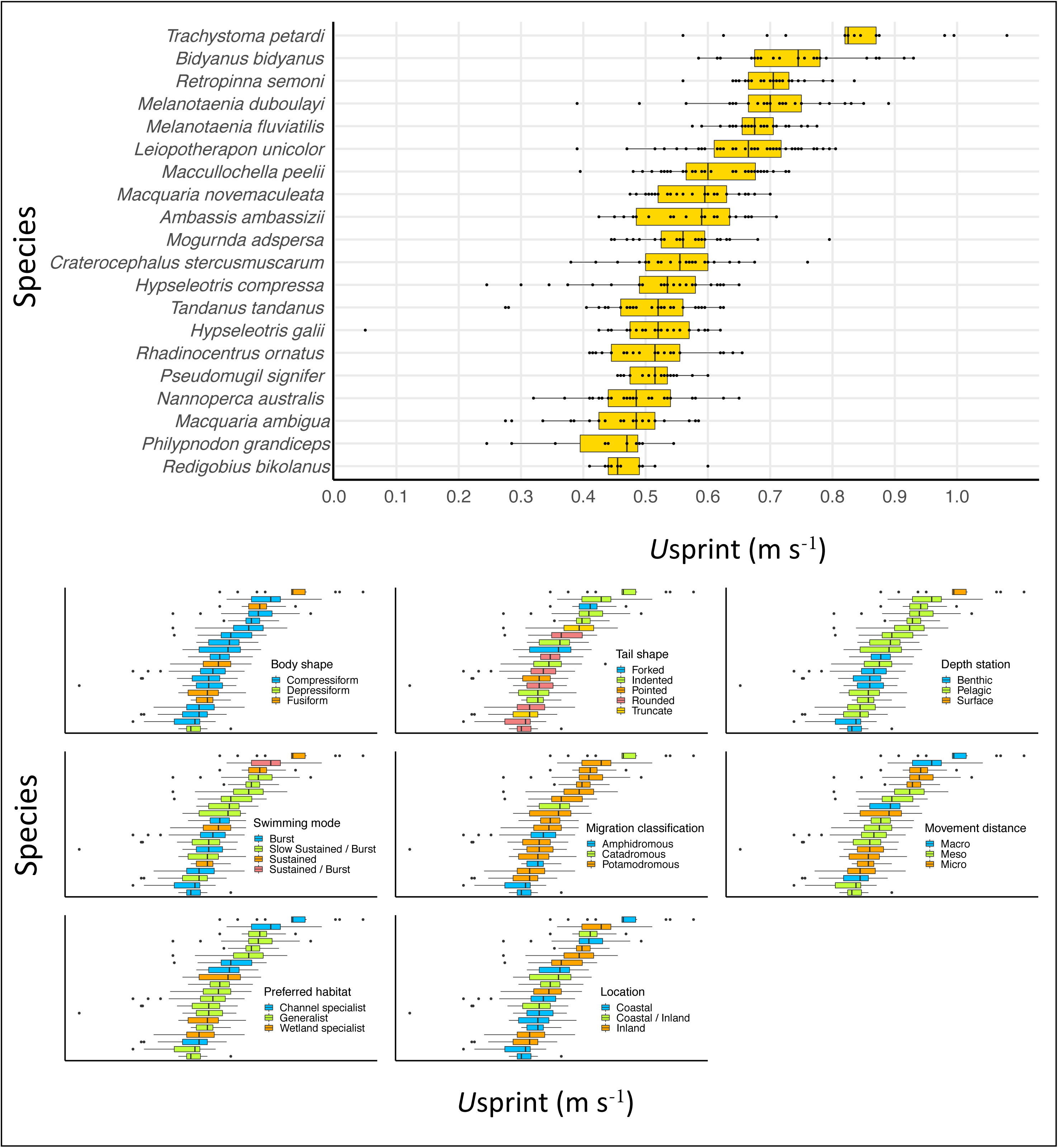
Visual comparison of the *U*sprint values with individual fish shown as dots. Each of the nine physical or behavioural characteristics is plotted to show the trends across the species tested.

### Characteristics – Ucrit (m s^−1^)

We modelled the predictor characteristics individually to assess their potential effect on the swimming performance of the species tested. See Fig. 1 for graphical presentation of comparisons but note that fish length was included in the modelling but not represented graphically. There was a statistically significant effect of depth station on the observed *U*crit values (F_2, 17.95_ = 9.5105, *p* < 0.002), with a post-hoc Tukey test revealing significant differences between all depth station comparisons (pelagic species performed better than benthic species (*p* < 0.005), surface swimming species performed better than benthic (*p* = 0.0002), surface performed better than pelagic (*p* < 0.008)). Body shape (F_3, 24.49_ = 3.173, *p* = 0.042) had a statistically significant effect on *U*crit, with fusiform species being better swimmers that compressiform species (Tukey; *p* = 0.033) (Fig. 1). Tail shape had a significant effect on *U*crit (F_4, 16.25_ = 5.6301, *p* < 0.005), with species with forked and indented tails performing better than those with rounded tails (*p* = 0.04 and *p* < 0.001). There was a statically significant effect of swimming mode on *U*crit (F_3, 17_ = 5.2459, *p* < 0.01), with a post-hoc Tukey test identifying significant differences with sustained swimming species performing better than burst swimming species (*p* < 0.002). Movement distance also showed a statistically significant effect on *U*crit (F_2, 18.21_ = 4.8085, *p* = 0.02), with species that perform macro-scale movements performing significantly better than those that perform meso-scale movement (Tukey; *p* = 0.01). Finally, migration classification also had a statistically significant effect on *U*crit (F_2, 18.15_ = 3.4201, *p* = 0.05), with catadromous species performing better than both amphidromous (*p* = 0.036) and potadromous species (*p* = 0.036). Location and preferred habitat had no significant effect on the observed *U*crit (m s^−1^) values.

### Usprint

#### Characteristics - Usprint (m s^−1^)

There was a statistically significant effect of tail shape on *U*sprint (F_4, 14.7_ = 4.4595, *p* = 0.014), with a post-hoc Tukey test revealing forked and indented tails were significantly better swimmers than pointed (*p* = 0.036, and *p* = 0.018). Indented tail species were also better performers than rounded tail species (*p* = 0.023) (Fig 2). As with *U*crit, there was a significant effect of depth station on *U*sprint (F_2, 17.70_ = 11.117, *p* = 0.0007), with the post-hoc Tukey test showing significant differences between all depth station comparisons; pelagic better swimmers than benthic (*p* = 0.002), surface better swimmers than benthic (*p* = 5.24e-05), surface swimmers better than pelagic (*p* = 0.003). Swimming mode had a significant effect on *U*sprint (F_3, 16.30_ = 3.4, *p* = 0.043), with the significant comparison being sustained swimming fish performing better than burst (Tukey; *p* = 0.0134). Migration classification also had a significant effect on *U*sprint (F_2, 17.64_ = 3.7098, *p* = 0.045). A post-hoc Tukey test revealed two comparisons with statistically significant differences, that being catadromous performed better than amphidromous (*p* = 0.0209) and potamodromous (*p* = 0.0442) species. Finally, there was no significant effect of body shape, location, preferred habitat or movement distance on *U*sprint.

### Predicting swimming performance (Ucrit)

Using all of the species characteristics to predict an unknown *U*crit decreased the accuracy and increased the variability of the predicted value. Fish length alone proved to be a robust characteristic for predicting a fishes swimming ability. Using only fish length, predicted *U*crit values were very similar to that of the observed *U*crit values for each species (Tukey post – hoc comparison *p* = 0.847). While this did not leave much room for improvement with the individual addition of either body shape, tail shape or depth station as a co-predictor with length, all of these co-factors gave an overall improvement to the predicted *U*crit values (all *p* = 1 meaning predicted values were overall very similar to the observed data). In most cases the observed data showed more variability than the predicted data, regardless of the characteristic(s) used (Fig. 3).

**Figure 3.**
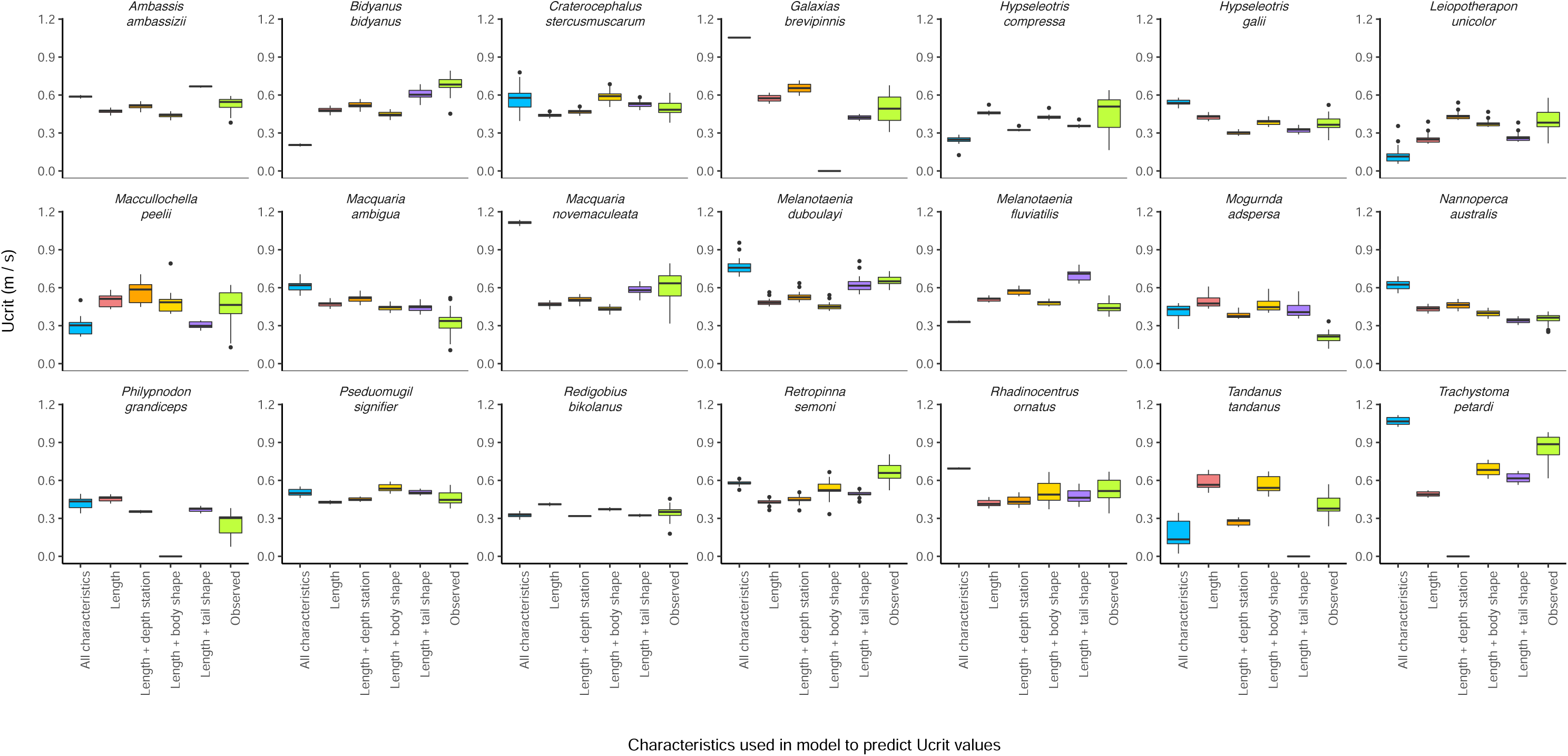
Comparison of observed values predicted *U*crit values. Five different predictive models were assessed using the nine physical and behavioural characteristics in Table 1.

### Modelled culvert traversability

The New South Wales (Australia) fish-friendly culvert guidelines recommend a maximum water velocity of 0.3 m s^−1^ during base flow periods. The culvert traversability modelling performed using the 25^th^ percentile *Ucrit* data suggests that this value is quite accurate as only four species, *M. adspersa, M. ambigua, P. grandiceps* and *R. bikolanus* would be unable to traverse the most common culvert length in NSW of 8 m (Fig. 4). The traversability modelling suggests that 75% the highest performing species, *T. petardi*, could traverse a 60 m culvert against a bulk channel flow of 0.6 m s^−1^.

**Figure 4.**
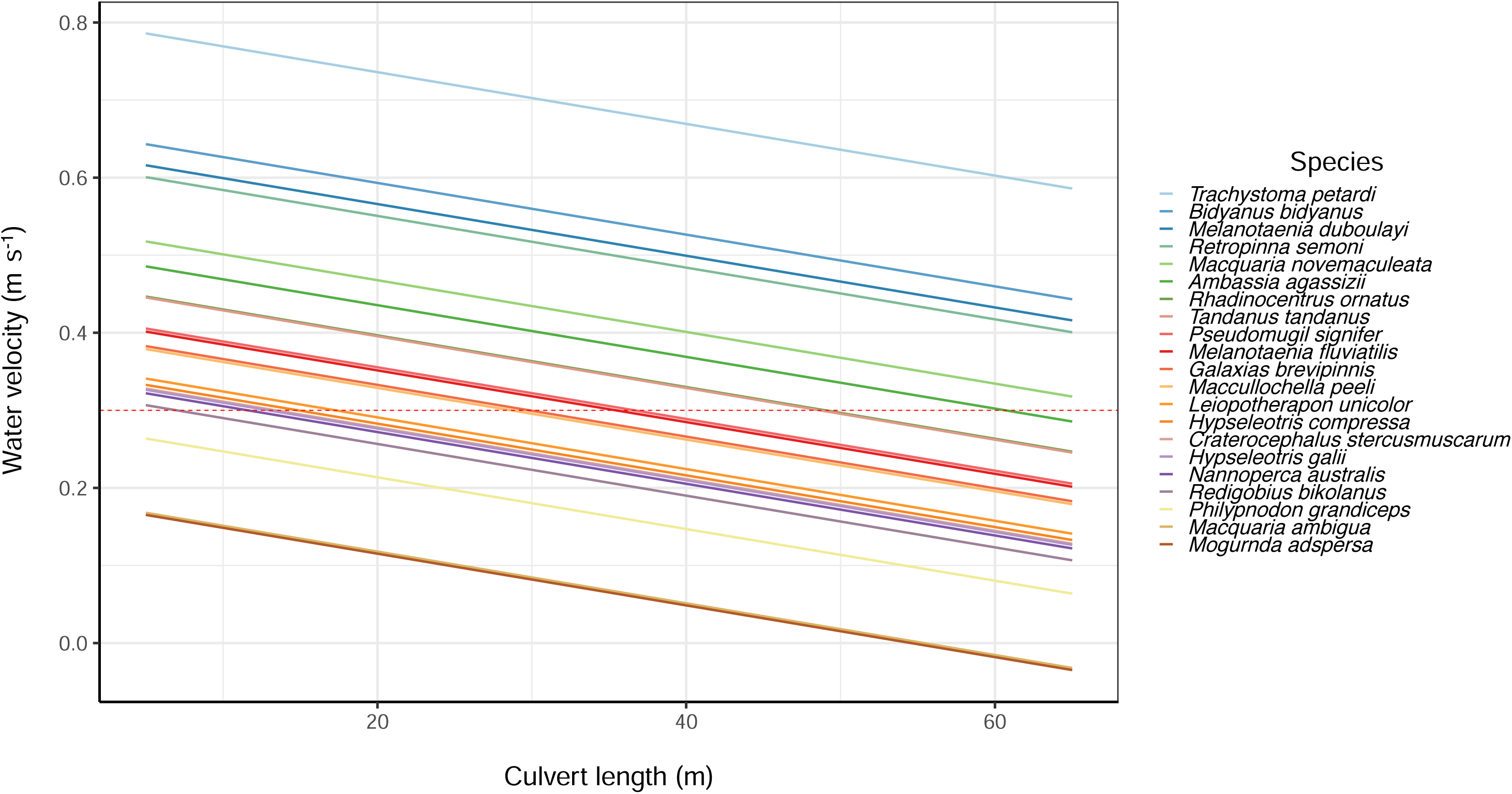
Culvert traversability modelling based on the 25^th^ percentile *U*crit value for twenty-two species of Australian native fish. By using the 25^th^ percentile value, the modelled curves show estimates where 75% of the population within the size range sampled could produce the required swimming performance to successfully traverse structures of different lengths. The red dashed line represents the recommended maximum water velocity (0.3 m s^−1^) during baseline flows in New South Wales, Australia.

### Swimming endurance and traverse success rates

The average swim times and size range of fish swum in the 12 m channel endurance trials are reported in Table 3. As expected, increasing the velocity decreased the average swimming endurance time for all species. The swimming endurance data was used to generate observed endurance curves for each species at each of the velocities it was swum at (Fig. 5).

**Figure 5.**
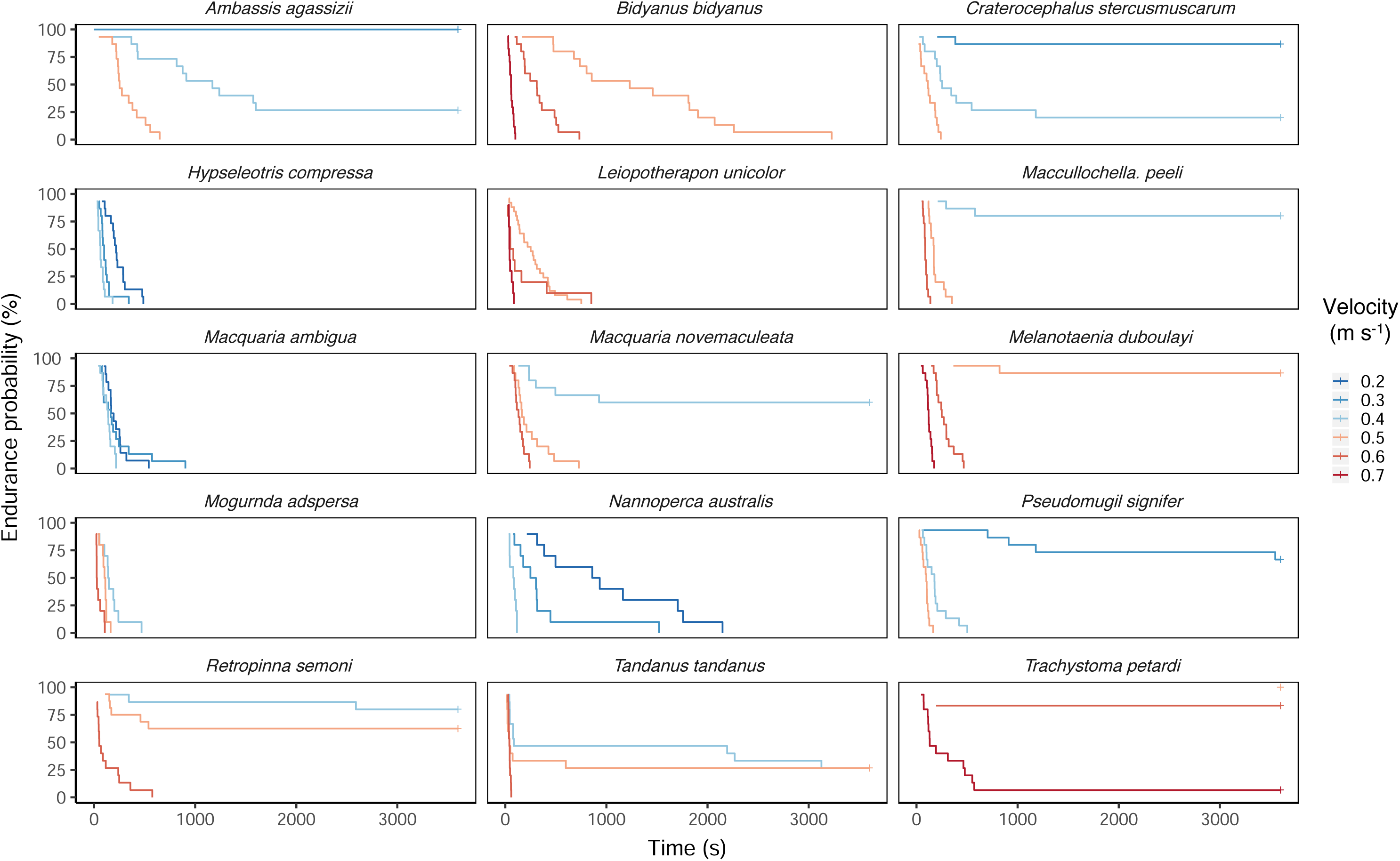
Endurance probability curves for fifteen species of Australian native fish. Each species was swum at three different test velocities reflective of their swimming capacity.

The trend of reduced swimming endurance times with increased velocity was repeated for the rates of traverse success, with increasing velocity reducing the percent of fish able to swim 8 m up the test channel without encouragement (Fig. 6). Modelling across all species showed a statically significant drop in the rates of traverse success between V1 and V2 (*p* = 0.0008), and between V2 and V3 (*p* = 0.0001). The exceptions to the overall trend, *M. duboulayi* and *T. tandanus*, can be explained by the size of the fish swum in the intermediate test velocity being slightly larger than that fish swum at the slower velocity. The average size of *M. duboulayi* was 8.6 cm in V1 and 9.8 cm in V2. Likewise, the average size of *T. tandanus* was 6.8 cm in V1 and 7.3 cm in V2.

**Figure 6.**
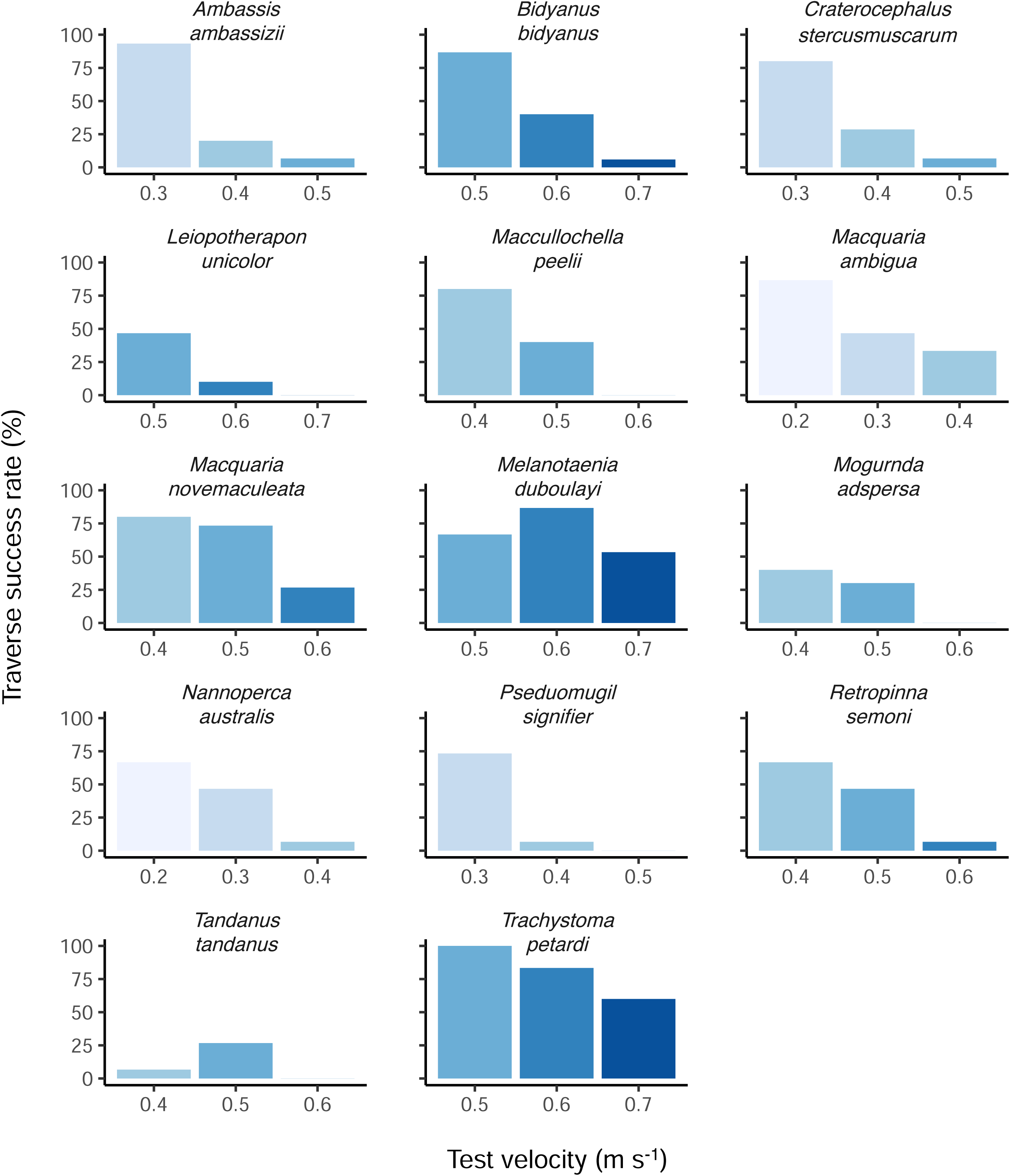
Rates of traverse success for fifteen species of Australian native fish. Each species was swum at three test velocities reflective of the swimming capabilities. Traverse success was defined as the individual fish swimming up 8 m of the experimental channel during the swimming endurance trial without encouragement.

## Discussion

Here we have reported an important baseline dataset characterising the swimming performance of twenty-one native Australian fish species. The purpose of this dataset is to enable fisheries managers and engineers to design appropriate fish-friendly instream structures (e.g. culverts and fishways), and to enable the remediation of existing problem sites based on quantified data. For these purposes it is important for policy makers to have high-quality scientific data to support and enforce their decisions.

An example are the guidelines for designing fish friendly road crossings in New South Wales, Australia, which currently recommend that baseline flows within culverts do not exceed 0.3 m s^−1^ (Fairfull and Witheridge, 2003). The determination of this velocity was based on limited swimming performance data for Australian freshwater fish. Using the traversable distance models generated from the average *U*crit data, we found that the 0.3 m s^−1^ recommendation is actually well supported, with all but two species unable to traverse 8 m, a distance equivalent to a culvert underneath a two-lane rural road crossing which predominate most waterway crossings (NSW DPI Fisheries unpublished data). This data indicates that there needs to be a contiguous region of the structure where water velocities do not exceed 0.3 m s^−1^ to facilitate the successful upstream passage of most Australian small-bodied fish.

While swimming performance is important information for designing fishways, obtaining detailed data on every species and size class is not always logistically feasible. Our results simplify this for Australian freshwater fish management through the diversity in morphologies and ecological traits covered in the species we sampled. This coverage allows the species we sampled to act as proxies for untested species that display similar physical attributes, such as body and tail shape, or behavioural traits, such as depth station, preferred habitat or movement distance. Interestingly the *U*crit data revealed that, with length as a cofactor, depth station was the best characteristic to infer the relative swimming performance of fish species for which no data exists. Our data suggest that in the context of Australian freshwater fish, body length followed by depth station, body shape and tail shape are the fundamental characteristics to infer the swimming capacity of un-tested fish species. Ultimately, the degree of difference in predictive power between these three characteristics was minimal, with fish length being the most important contributing factor. These results may allow fisheries managers in Australia to use the existing data we present to infer the likely swimming performance for a species of interest of given body length for which performance data is lacking, especially if a secondary characteristic – depth station, body or tail shape – is taken into account. As fish size was highly correlated with swimming performance, extrapolating performance estimates for individuals outside the size range we tested is also feasible.

It is important to acknowledge certain limitations to the utility of this dataset for broadly categorising general fish swimming performance, or for making sweeping assumptions of a tested species’ performance. First, the fish we tested did not cover all size classes that may be of interest for the management of a particular species at a location of interest. We focussed on small fish as they face a greater velocity challenge than larger fish. Secondly, logistical constraints meant we could not use wild-caught individuals or collect swimming performance data for all ontogenetic stages of the species tested. The use of captive bred fish (exceptions being *R. semoni, M. duboulayi, P. signifier, P. grandiceps* and *R. bikolanus*) means that our reported data is possibly conservative, as other work has shown the swimming performance of wild-caught fish can exceed that of captive bred (Basaran et al., 2007).

Thirdly, many species including gudgeons, galaxiids, salmon parr and darter species, employ specialised swimming or movement techniques, such as the use of pectoral fins to climb or station hold (Carlson and Lauder, 2011; Peake et al., 1997; Webb, 1989). During the *U*crit and *U*sprint trials, *H. compressa, H. galii, M. adspersa, P. grandiceps* and *R. bikolanus* all displayed station holding behaviour, particularly when being swum at velocities that were not challenging. This was again observed by *M. adspersa* during endurance trials in the 12-meter experimental channel. At challenging velocities, these species were unable to utilise station holding due to losing their grip on the smooth PVC surfaces in the experimental equipment. Man-made in-stream structures are typically made with concrete that has a rougher surface, meaning it is reasonable to anticipate that climbing and station holding species may be capable of withstanding faster water velocities in a real situation than what was observed in the experimental flumes. Furthermore, a fish’s motivation to perform is hard to quantify both in the field and laboratory (Goerig & Castro-Santos, 2017; Goerig et al., 2019). The use of volitional swim tests conducted in real or simulated open channels has increased in the literature since 1990 and should be conducted where possible to support data collected in chambered flumes where physical constraints may impact results. The endurance and traverse success data collected for fifteen species in a 12 m channel can be used to support management decisions based on the reported *U*crit and *U*sprint data.

Finally, fish swimming performance is greatly affected by temperature. We standardised our dataset by holding the fish and quantifying their swimming performance at 25°C. While this temperature generally approximates a late spring or summer temperature in most coastal Australian rivers, it may be outside the thermal optima for some of the more temperature species that we tested. Natural waterbodies have significant seasonal, daily and spatial variations in temperature that directly affect the ability of a fish to swim (Caissie, 2006).

We focussed on reporting our data in m s^−1^ as it is directly relevant to fisheries management as it more closely reflects commonly reported water velocity units. While fish swimming performance is often reported and discussed in body lengths per second for comparative purposes, this is not applicable to real world situations faced by water infrastructure engineers and fisheries managers. Simply, in a real-world situation fish are facing the same velocity challenge, and a small fish doing ten body lengths per second faces a greater challenge than a much larger fish capable of five body lengths per second. The physical size difference also means that small and large fish face different hydrological conditions, and resulting challenges, when in the exact same location within both the natural environment and an artificial in-stream structure. The size difference means alternate energy saving strategies may be employed, with small-bodied fish able to utilise boundary layers to reduce the velocity challenges found within structures such as culverts (Goodrich et al., 2018; Watson et al., 2018). While excessive turbulence can disorientate and exhaust small-bodied fish, larger growing species may use turbulence to propel themselves forward if the eddies match their body size (Liao et al., 2003a,b; Taguchi and Liao, 2011).

Our results have highlighted that while *U*sprint is an informative metric for fisheries management, it does not approximate the sprinting capabilities of fish, which is defined as the maximum sustainable effort up to 20 s. Most of the *U*sprint trials had fish swimming for around 1 - 2 min before fatigue, with each species’ average values not much greater than the *U*crit average. This is a limitation of the experimental design used by Starrs et al. (2011), where the progressive increase in velocity exhausts the fish before the max burst swimming velocity is obtained. For comparison, Mallen-Cooper (1992) measured the swimming performance of three sizes of Australian Bass through a vertical slot (around 200 mm sprint distance) and determined velocities at which 95% of bass were predicted to pass through the slot. The determinations were 1.02 m s^−1^ for 40 mm bass, 1.40 m s^−1^ for 64 mm bass, and 1.84 m s^−1^ for 93 mm bass, noting that these are water velocities through the slot, not the velocities that the fish are required to achieve forward progress. These values are considerably greater than the highest *U*sprint recorded for bass in the current trials (∼0.7 m s^−1^). We agree that what Mallen-Cooper (1992) tested is much closer to the max burst speed than what *U*sprint quantifies. Instead, we see *U*sprint sitting between a prolonged swimming speed, and a fish’s burst speed, and we actually see great utility in it as a double check of proposed velocities through certain instream infrastructure where a fish would need to traverse any distance greater than a few hundred millimetres (e.g. a vertical slot fishway), but less than a few metres. A culvert that would have velocities closer to the fish’s *U*sprint is highly unlikely to pass that species, and thus becomes an upper limit for any considerations of water velocities and swim distances greater than 500 mm. We suggest that *U*sprint should be either renamed, or redefined, as we are not confident it represents the burst capabilities of the fish. The cost implications of using it in proposing fishway velocity design criteria where burst swimming is required (e.g. through vertical slots) would be significant, as even conservative fishway designs have burst velocity criteria well over 1.0 m/s, with most at 1.4 m/s and some up to 1.8 m/s depending on the target fish.

The utility of laboratory-defined fish swimming performance data for the design of fish passage structures has been in wide usage throughout much of the world (Botha et al., 2018; L Cai et al., 2018; Scruton et al., 1998; Katopodis et al., 2019; Link et al., 2017; Peake, 2014; Starrs et al., 2011; Bice and Zampatti, 2005). However, a similar resource did not exist for Australian freshwater fish species meaning that many fish passage structures were inappropriate for their passage requirements. We have collected the first comprehensive database of fish swimming performance capacities for Australian fish. From this data we have been able to identify key morphometric and behavioural traits that can be used to predict swimming performance in similar species for which performance data does not exist. Moreover, these data focus on the most vulnerable size class of fish in the environment, small individuals, less than 10 cm TL. The data presented here represents a significant resource for fisheries managers and water infrastructure engineers in Australia to inform and develop more species-specific ‘fish-friendly’ fish passage in Australian waterways and to promote passage in a wider range of body shapes and sizes than before.

## Supplementary information

### Results

#### Characteristics - Ucrit (BL s^−1^)

Overall, the groupings of species with the same characteristic seemed clearer with *U*crit presented in body lengths per seconds (Fig. 1 and Fig S1), but the overlap between species increased, reducing the statistical significance of comparisons. The only statistically significant effect was of body shape on *Ucrit* BL s^−1^ (F_3, 24.342_ = 5.5199, p < 0.005), with the post-hoc Tukey test showing fusiform species to be significantly better swimmers than species with either compressiform (*p* = 0.006), depressiform (*p* = 0.042), and sagitiform species (*p* = 0.042) (Fig. S1). Tail shape, swimming mode, depth station, location, preferred habitat, migration classification, and movement distance had no statistically significant effect on *Ucrit* BL s^−1^.

#### Characteristics - Usprint (BL s^−1^)

The only statistically significant effect on *U*sprint (BL s^−1^) was by body shape (F_2, 17.277 = 3.8772_, p = 0.04). A post-hoc Tukey test revealed that the only significant difference was with fusiform performing better than compressiform body types (*p* = 0.0274). There was no significant effect of tail shape, swimming mode, depth station, preferred habitat, location, migration classification or movement distance on *U*sprint (BL s^−1^) (Fig. S2).

#### Figure captions

**Figure S1.**
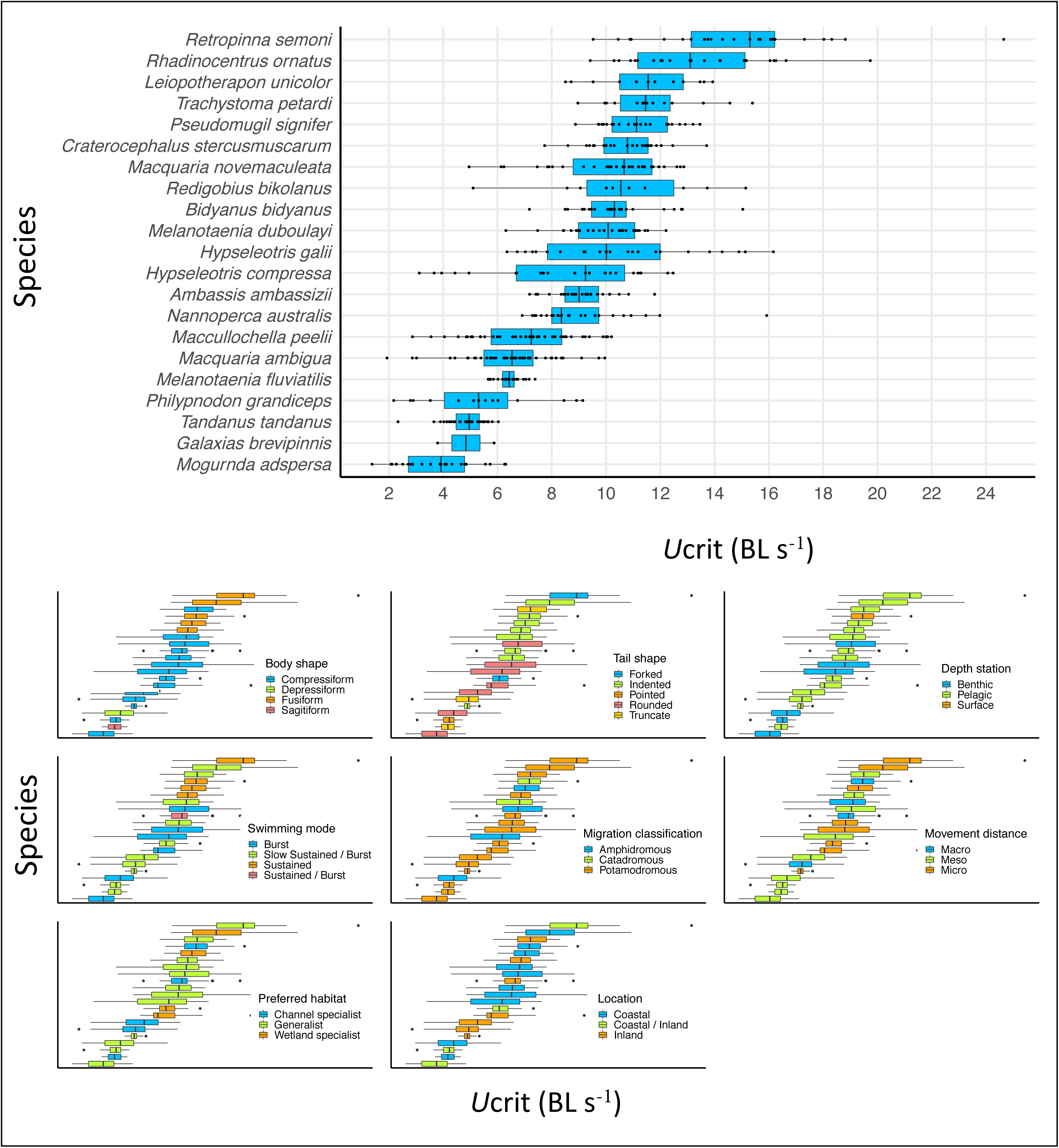
Visual comparison of the *U*crit (BL s^−1^) values with individual fish shown as dots. Each of the eight physical or behavioural characteristics are plotted below to show the trends across the species tested.

**Figure S2.**
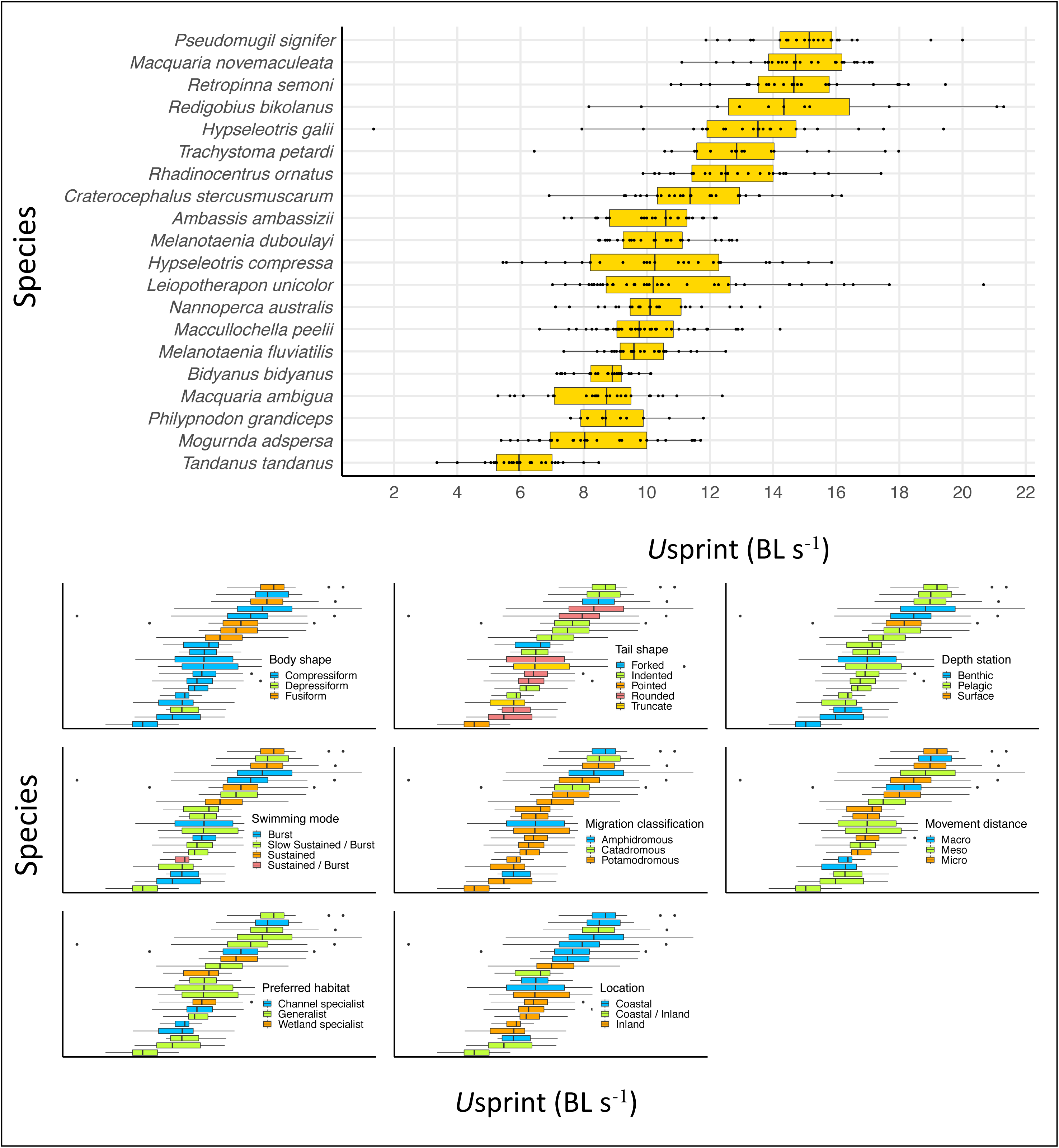
Visual comparison of the *U*sprint (BL s^−1^) values with individual fish shown as dots. Each of the eight physical or behavioural characteristics are plotted below to show the trends across the species tested.

